# Towards an Open Data Framework for Body Sensor Networks Supporting Bluetooth Low Energy

**DOI:** 10.1101/076166

**Authors:** Ninoshka K. Singh, Darrell O Ricke

**Author notes:** This work is sponsored by the Assistant Secretary of Defense for Research & Engineering under Air Force Contract #FA8721-05-C-0002. Opinions, interpretations, recommendations and conclusions are those of the author and are not necessarily endorsed by the United States Government.

## Abstract

Major companies, healthcare professionals, the military, and other scientists and innovators are now sensing that fitness and health data from wearable biosensors will likely provide new discoveries and insights into physiological, cognitive, and emotional health status of an individual. Having the ability to collect, process, and correlate data simultaneously from a set of heterogonous biosensor sources may be a key factor in informing the development of new technologies for reducing health risks, improving health status, and possibly preventing and predicting disease. The challenge in achieving this is getting easy access to heterogeneous data from a set of disparate sensors in a single, integrated wearable monitoring system. Often times, the data recorded by commercial biosensing devices are contained within each manufacturer’s proprietary platform. Summary data is available for some devices as free downloads or included only in annual premium memberships. Access to raw measurements is generally unavailable, especially from a custom developed application that may include prototype biosensors. In this paper, we explore key ideas on how to leverage the design features of Bluetooth Low Energy to ease the integration of disparate biosensors at the sensor communication layer. This component is intended to fit into a larger, multi-layered, open data framework that can provide additional data management and analytics capabilities for consumers and scientists alike at all the layers of a data access model which is typically employed in a body sensor network system.

## I. INTRODUCTION

As stated by [4], one of the main goals of mobile health frameworks is to foster the research and development of health, fitness, and other medical studies as well as to accelerate the development of mobile health technologies and applications.

The Integrated Biomedical System (iBio) study collects a diverse set of physiological health data from multiple heterogeneous, wearable sensors [8]. The data streams for each commercial off-the-shelf (COTS) wearable sensor are unique, if available at all. Sensors compliant with the Bluetooth Low Energy (BLE) communications standard provide the easiest access to raw data across multiple COTS sensors supporting BLE. Thus, the study focused on collecting data from sensors supporting BLE. This paper proposes an open data framework for collecting raw data from multiple wearable sensors using a smart mobile device to cache and forward sensor data to remote storage sources. The major implementation challenge faced was trying to simultaneously access and interpret a variety of physiological data from a collection of disparate biosensors produced by different manufacturers.

Most wearable mobile health technologies provide a common set of data access capabilities in a multi-layered access model shown in Fig. 1. Today, in industry, this model is typically implemented as a closed, proprietary technology platform by different companies who are either producing a wearable monitoring device (e.g. Hexoskin’s smart shirt [11], Moticon’s smart foot insoles [14]) or a collection of wearable devices as a part of an ecosystem [12, 13]. The typical flow of data in this model involves the wearable monitoring system periodically transmitting data over one or more ultra-low power wireless channels to a central aggregation device such as a smart mobile phone. Transitory live data can then be viewed on the mobile device’s display using a mobile app or it can be persisted locally if the app has access to the raw data. Typically, having raw data access is only possible through a custom developed mobile app using a software API [11] which is generally provided at a price set by the company. Additionally, data can also be pushed to remote storage over a higher speed wireless link for later viewing, usually only as summary statistics on the web. Furthermore, some companies allow data to be downloaded post-activity, for a fee. Sometimes tools are also provided for the user to analyze their data in an intuitive way offline or through the mobile app. Again, usually the data analytics tools of a company’s platform require the user to pay a price, such as Moticon’s Beaker software [14].

**Figure 1.**
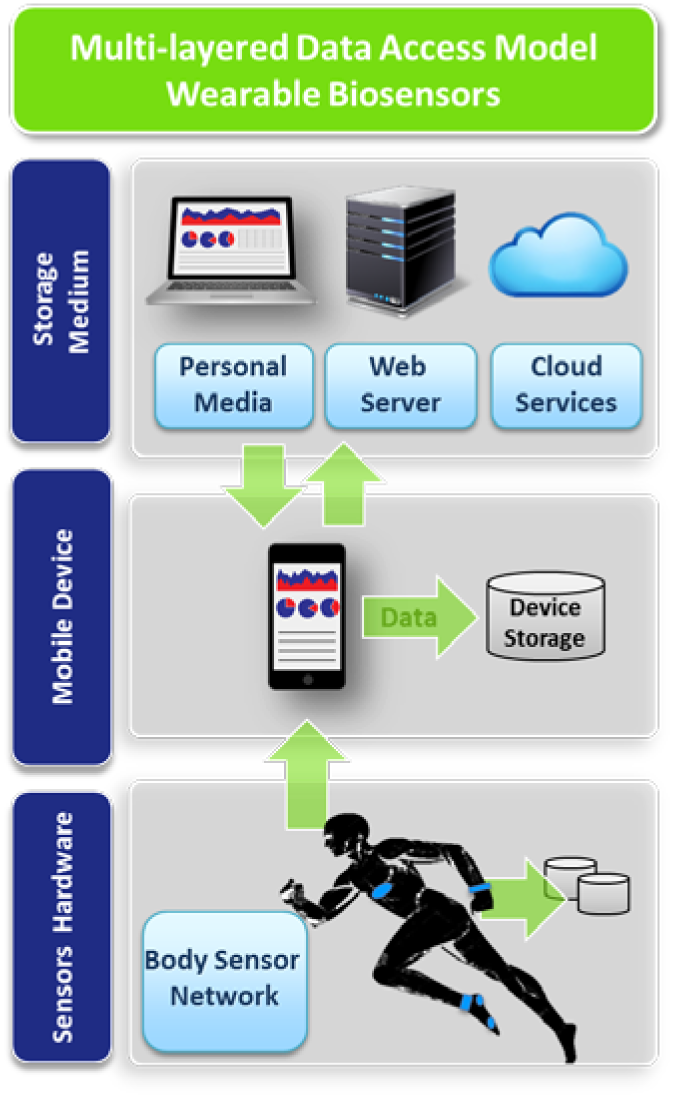
Diagram illustrates how data moves wirelessly and where it is stored and accessed, in a multi-layered data model which is typically implemented for BSN systems.

All of these restrictions in commercial technology platforms severely limit how consumers, healthcare professionals, military, scientists, and other innovators can use the health and fitness data. Arguably, the most significant source that contributes to these data access restrictions is in how the underlying communication is implemented between the sensor network and the central data aggregation device for data exchange. Even though many companies support BLE in their products, they generally only implement the lowest layers of the communication protocol and then build proprietary messaging on top of this to encapsulate and transmit their data to a central mobile device. Few products take advantage of BLE’s standardized services and profiles feature for health data encapsulation, exchange, and formatting. Having a sensor communication layer which takes advantage of efficient, standardized data exchange and formatting is paramount for improving interoperability across biosensors and for promoting a unified data model in an open data framework (shown in Figure 2).

**Figure 2.**
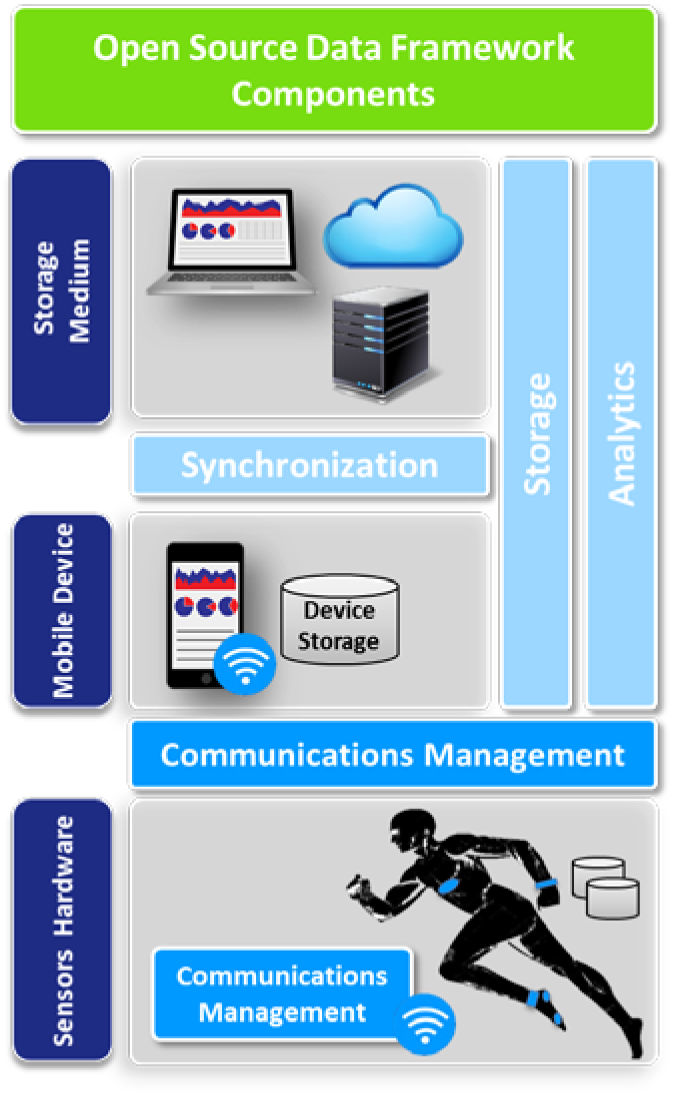
The blue blocks illustrate the components of an open data framework. The darker blue components represent the BLE communication integration components proposed and discussed in this paper.

The remainder of this paper is organized as follows. Section II analyzes the design of Bluetooth Low Energy and discusses why it is ideally suited to satisfy open data access and interoperability goals needed at the sensor communication level. Furthermore, this section expands on how the Host Controller Interface and Generic Attribute Profile (GATT) layers [3] can each be abstracted or extended to support these goals. Section III discusses the design and implementation of a multi-sensor data collection mobile app, and how parts of it can be abstracted into the open data framework’s communication components discussed in Section II. Finally, Section IV discusses the future work required to implement all components of the open data framework.

## II. BLUETOOTH LOW ENERGY

The advent of Bluetooth Low Energy (BLE) has occurred while other low-power wireless solutions, such as Zigbee, 6LoWPAN or Z-Wave, have been steadily gaining momentum in application domains that require multi-hop networking. However, BLE constitutes a single-hop solution which is especially applicable to use cases such as in healthcare, sports, and general fitness applications [2]. From an energy efficiency perspective, BLE is particularly advantageous for its ultra-low power consumption, which is an important requirement for both consumer and military real-time physiological monitoring applications [2]. BLE optimizes for this by using low duty cycle transmissions, very short transmission bursts between long periods of inactivity, and extremely low power sleep modes [1, 2]. From a data access perspective, BLE separates the data communication protocol from the data structure, format, and behavior definitions. This provides simple, reusable, and extensible data encapsulation and exchange. The combination of BLE’s flexible and extensible data access features with its low power design make it a powerful candidate for body sensor network applications. The focus of this section will be on understanding some of the data access design features and how they can be extended to enable a mobile device to easily collect data from a single network composed of wearable heterogeneous data sources which could be commercial or prototype based.

### A. Architecture

The architecture of BLE is split into three basic parts: controller, host, and applications (see Figure 3). The controller is the lowest level firmware/software that runs on the radio transceiver hardware. The controller layers include the Physical and Link layers, and the lower part of the Host Controller interface (HCI). These layers define how a physical device can transmit and receive bits or bytes of information using radio signals and how a device connects to another device. The host layers, which include the upper part of the HCI, define higher level protocols, which specify how two or more devices exchange and access data. The applications use the host and controller software layers to enable different use cases [3].

**Figure 3.**
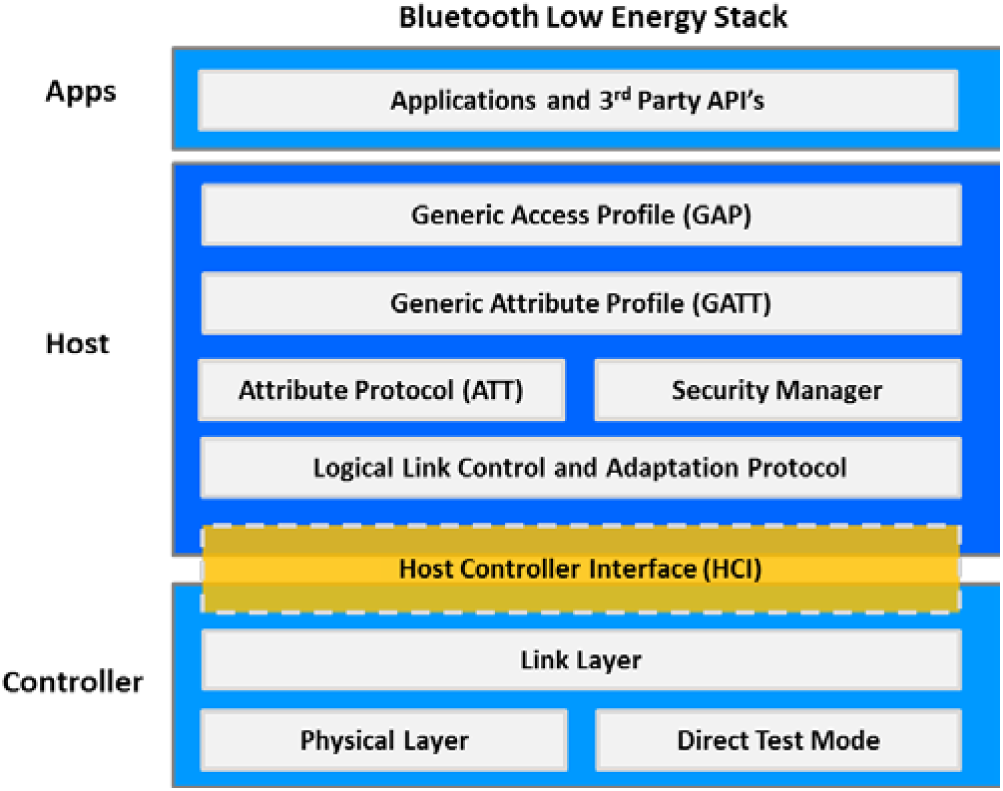
Illustration of Bluetooth Low Energy Stack for applications (apps) to communicate with the host device (e.g., Android phone) interacting with the wearable device via the Controller.

### B. Object-Oriented Design Features

If the host layers are used, all application data is exchanged and encapsulated in units of information called *attributes* [3]. An attribute is a piece of labeled, addressable data, composed of a unique identifier, the type, and a value that defines the actual state data that’s encapsulated. This data is organized hierarchically in sections called *services*, which encapsulate conceptually related pieces of user data into *characteristics*. A characteristic can also expose behavior [3]. Each service contains one or more characteristics and can reference or extend other services. Because of this atomic design, services can be reused by multiple clients and can be composed together to create more complex services. Finally, a *Profile* [3] is a specification used by a client device that defines a set of rules for discovering, connecting, configuring, and using services.

### C. Attribute Protocol and Generic Attribute Profile

The Attribute Protocol (ATT) [3] is the data communication protocol by which a client can find and access attributes on a device containing an attribute server. The Generic Attribute Profile (GATT) is used on top of the Attribute Protocol. This specification defines additional procedures that define standard ways for services, characteristics, and their descriptors to be discovered and then used. A more generic client that wants to dynamically access pre-existing or new services as they become available can use a procedure to enumerate the entire attribute database. Furthermore, if the characteristics include descriptors that allow them to be self-describing, the client can dynamically parse the information without having any prior knowledge about the data.

## III. FRAMEWORK APPROACH

In the previous section, an overview of the BLE architecture and its key design features was analyzed. In this section, the BLE communication components of the open data framework are discussed. These components aim to generalize biometric data exchange over the higher level ATT/GATT protocols and over the lower level Host Controller Interface. The goal is to provide data exchange such that both commercial and prototype sensors can be accessed in a uniform way on a data aggregation device.

### A. Supporting Standard and Generic GATT Services

Profiles and services provide a modular architecture, which innovators can reuse and extend. Both commercial and prototype sensors can leverage this architecture to provide access to data in a standard way.

An advantage of the GATT specification is that it allows clients to discover services in different ways. Typically, a simple client with limited resources may only care about finding a single certified Bluetooth SIG service such as the Heart Rate Service [9], which it knows how to parse. However, a more generic client such as a mobile device connected to a network of heterogeneous sensors, may need to dynamically access both known certified services, and new, arbitrary services containing non-standardized health data measured by prototype sensors.

To support this case, a new self-describing service is needed. In Figure 4, a *Generic Biometric Service* definition is proposed. This service contains a single measurement characteristic that contains a multi-byte value field, along with four descriptors [3] describing exactly how a generic client can parse the value field and understand it in a human readable form. The first descriptor is required for the client to receive updates when the value changes. The User Description descriptor allows a human readable text string to be associated with the characteristic. The Presentation Format Descriptor contains metadata describing the format, exponent, and units of the measured data. Additionally, if the data contains related, but separate parts, the Aggregate Format Descriptor can be used which allows the service to specify a separate Presentation Format Descriptor (PFD) for each part. The Aggregate Format Descriptor specifies which PFD is used for each part [3].

**Figure 4.**
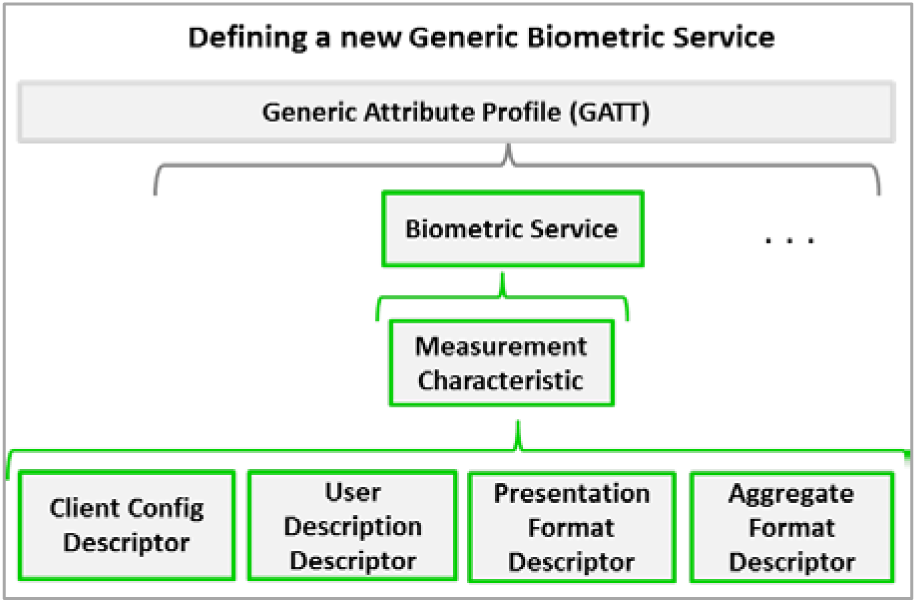
Proposed Generic Biometric Service with a Measurement Characteristic composed of four descriptors that make the characteristic self-describing.

### B. Open Messaging Protocol over Host Controller Interface

For many devices, a Host/Controller Interface (HCI) firmware will be provided which allows a host that implements the full BLE stack, to communicate with a radio chip that only implements the controller part of the stack. BLE accounts for this asymmetry by separating communication behavior into different roles, which implies that the most resource-constrained devices can implement minimum processing and control logic while a more computationally able host (e.g. mobile device) can implement more functionality required to manage connections to devices, collect data, and process it.

This architectural split has been extremely popular in Bluetooth classic, for which over 60 percent of all Bluetooth controllers are used through the HCI interface [2]. Many BLE radio chip manufacturers provide a simple HCI application programming interface (API) for developers to use. However, the main limitation is that the API’s are proprietary and often not universal across radios. In Table I, the message format for data exchange over HCI is proposed. An open messaging protocol over HCI will allow all developers to interact with BLE controllers in a unified way regardless of the APIs that come with vendor-specific, commercially available BLE radios. These messages can encapsulate both command information and/or data. A future paper will discuss more details on this protocol.

**TABLE I.**
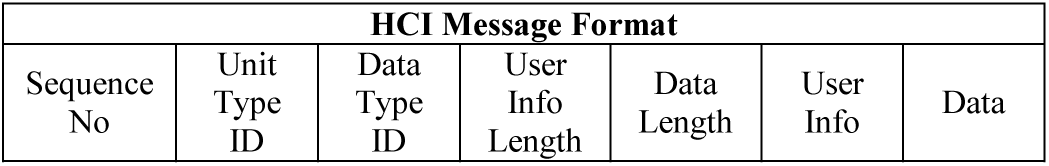
MESSAGE FORMAT FOR OPEN MESSAGING PROTOCOL OVER HCI

| HCI Message Format |  |  |  |  |  |  |
| --- | --- | --- | --- | --- | --- | --- |
| Sequence No | Unit Type ID | Data Type ID | User Info Length | Data Length | User Info | Data |

## IV. MOBILE APPLICATION FOR MULTI-SENSOR DATA COLLECTION

An Android application was developed which aggregates and records raw data from multiple commercial biosensors. The goal was to be able to periodically push the data to a remote storage service developed by MIT Lincoln Laboratory (MITLL), called Integrated Biomedical System. MITLL’s data system provides open source, data management and analytics capabilities for physiological, genomic, interactome, and exposome data sources. The Integrated Biomedical System description and written consent form was reviewed and approved by the MIT (IRB) Committee on the Use of Humans as Experimental Subjects (COUHES) for volunteers.

A number of commercial biosensors were considered for use in the multi-sensor data collection task. Out of the devices shown in Table II, the Mio Link, Polar H6 monitor, Spree headband, Jabra Sport Pulse earphones, and Scosche Rhythm+ wristband all provided access to raw biometric data over Bluetooth Low Energy, using standard certified health services discussed in Section II. These sensors were used in the data collection mobile app. The Zephyr BioHarness3, Hexoskin, and SenseCore smart shirts only used Bluetooth Classic to transmit data. These devices most likely implemented their own proprietary data protocol on top of the Serial Port Profile [10] to transmit data. Thus, it was not possible to receive data in the app directly. The Basis Peak, Empatica E3, and Microsoft Band sensors operated using BLE but did not provide access to the raw data for free. Therefore, these sensors were not used in the app.

**TABLE II.** EVALUATED COMMERCIAL BIOSENSORS FOR DATA COLLECTION APP DEVELOPMENT

| Product | Company | Bluetooth Protocol | Biometrics | Direct access to Raw Data | Mobile OS Support |
| --- | --- | --- | --- | --- | --- |
| Mio Link (Wrist) | Mio Global | Low Energy | Heart Rate | Yes, via custom App | Android<br>iOS |
| Spree (Head) | Spree | Low Energy | Heart Rate<br>Skin Temperature | Yes, via custom App | Android<br>iOS |
| Polar H6 (Chest) | Polar | Low Energy | Heart Rate | Yes, via custom App | Android<br>iOS |
| Jabra Sport Pulse (Ear) | Jabra | Dual Mode: Classic/Low Energy | Heart Rate | Yes, via custom App | Android<br>iOS |
| LifeBeam Hat | Scosche Industries | Low Energy | Heart Rate | Yes, via custom app | Android<br>iOS |
| Rhythm+ (Arm) | Scosche Industries | Low Energy | Heart Rate | Yes, via custom App | Android<br>iOS |
| BioHarness 3 (Chest) | Zephyr | Classic | Heart Rate, Breathing Rate, Posture, GPS, Activity | No, requires API access | Android<br>iOS |
| Hexoskin Smart Shirt | Hexoskin | Classic | Heart Rate, Breathing volume, HR Variability, Breathing rate, Activity, Sleep | No, requires API access with paid subscription | Android<br>iOS |
| Basis Peak (Wrist) | Intel | Low Energy | Heart rate, activity, sleep | No, requires API access with paid subscription | Android<br>iOS |
| E3 (Wrist) | Empatica | Low Energy | PPG, EDA, Temp Activity | No, requires API access with paid subscription | Android<br>iOS |
| Microsoft Band (Wrist) | Microsoft | Low Energy | Heart Rate, | No, only summary statistics | Android<br>iOS |
| SenseCore Smart Shirt | SenseCore AG | Classic | Heart Rate, Breathing rate, Activity, Sleep | No, only summary statistics | Android<br>iOS |

### A. Android Application Architecture

**Figure 5.**
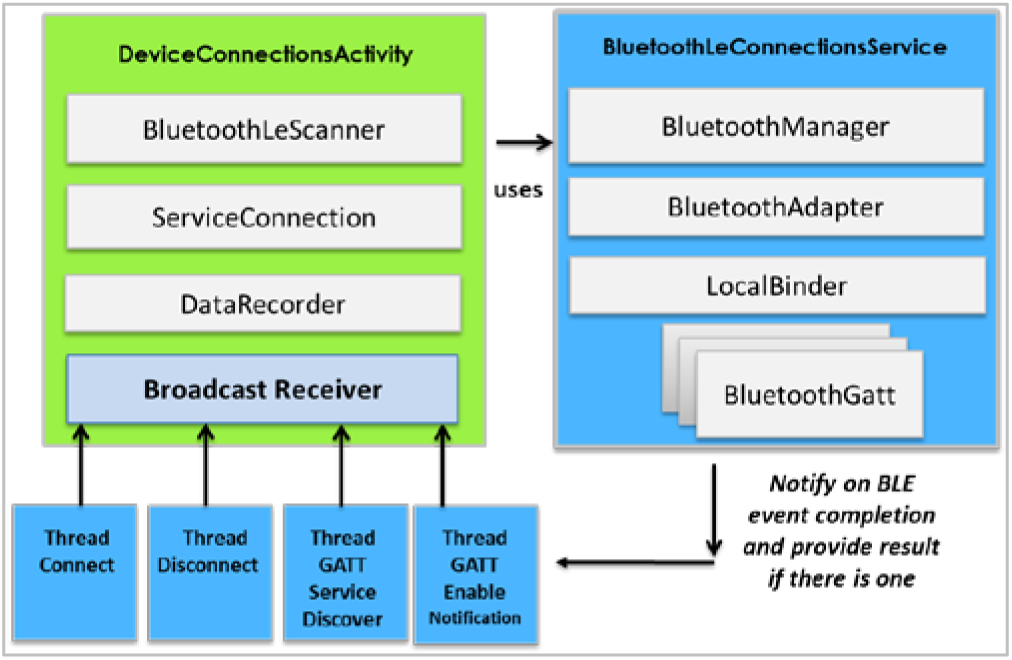
Application architecture for communicating with multiple BLE biosensors on a mobile device using Android’s BT/BLE API [15].

In Figure 9 the architecture of the implemented Android application is shown. The application is composed of three Android framework components: an activity, broadcast receiver, and a service. The *DeviceConnectionsActivity* is responsible for rendering UI updates and invoking other components to carry out BLE operations such as device discovery, connections management, and data collection from multiple BLE sensing devices. Additionally, this activity has a reference to a Broadcast Receiver, which is responsible for receiving asynchronous notifications from the *BluetoothLeConnectionsService* whenever a GATT action occurred or is completed (e.g. Service Discovery, Characteristic notification, update from a sensor, etc.…). The *BluetoothLeConnectionsService* manages connections to multiple sensor devices in separate threads. Once a connection is established with a sensor device, the application invokes all data exchange in the corresponding thread. This includes the GATT procedures for discovering what services are present on the device’s GATT server and the procedure for enabling automatic notifications when data is updated within that service. In the current implementation, the data parsing for specific, certified SIG services such as Heart Rate and Health Thermometer are hard-coded into the application. Ideally, this should be dynamic. The application could leverage a framework component, which does service registration. The user can select the desired biometric properties they want to measure, and the application can call upon a services registry component which can map the selected properties to certified services that may contain the corresponding characteristics for these data types. Finally, the application can invoke standard routines for parsing data delivered by any of these registered services (i.e. published on Bluetooth SIG website as open standards).

### B. Application Data Collection Results

To demonstrate the practical use of the app, the Mio Link wristband, Polar H6 chest monitor, Spree headband, and Scosche Rhythm+ wristband were worn simultaneously to collect data on a Samsung Galaxy S5 smartphone. The heart rate (HR) data results for two separate experiments that were 12.5 minutes and 7.5 minutes in duration, respectively, are shown in Figures 6 and 7. A few different exercises were conducted to illustrate the various HR ranges. It can be noted that the Polar H6, Mio Link, and Rhythm+ were all fairly well correlated. The fit of the Spree headband appears to have not been optimal. Further experiments would need to be conducted to verify this.

**Figure 6.**
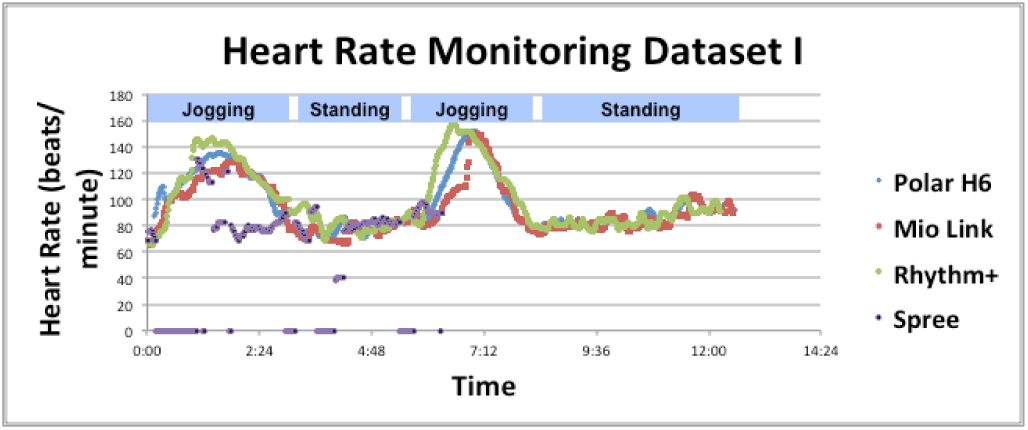
Shows a simultaneous heart monitoring from four biosensors while performing a series of activities.

**Figure 7.**
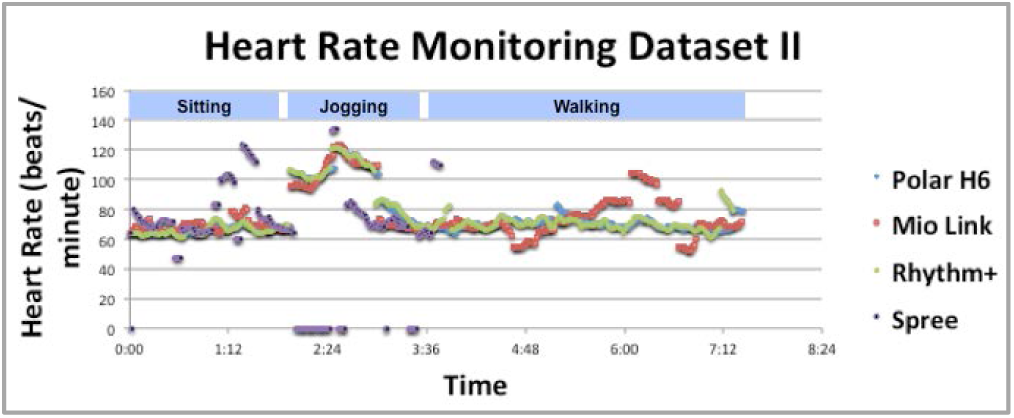
Shows the results of a second data collection which shows simultaneous heart monitoring from four biosensors while performing a different combination of activities.

### C. Abstracting the Application into a Client-Server Framework Architecture

Most mobile platforms (i.e. Android and iOS) provide basic Bluetooth Low Energy API’s that provide abstractions to the native software stack running on a vendor-specific mobile device. However, as was discovered from application development, typically, these API’s do not implement the necessary, more complex client-side functionality that requires the application to manage communications with multiple BLE devices simultaneously. Furthermore, the application must implement ways of processing different types of GATT service data contained on the sensing devices. Similarly, at the Host Controller Interface, the application is responsible for knowing how to interpret different, proprietary sets of commands from various sensors using vendor-specific BLE radio chips.

For each communication mode, Figure 8 and 9 present an overview of the architecture for both the client-side (e.g. mobile device) and server-side (e.g. biosensor device) software components required to abstract the higher level communication over ATT/GATT and the lower level Host Controller Interface.

**Figure 8.**
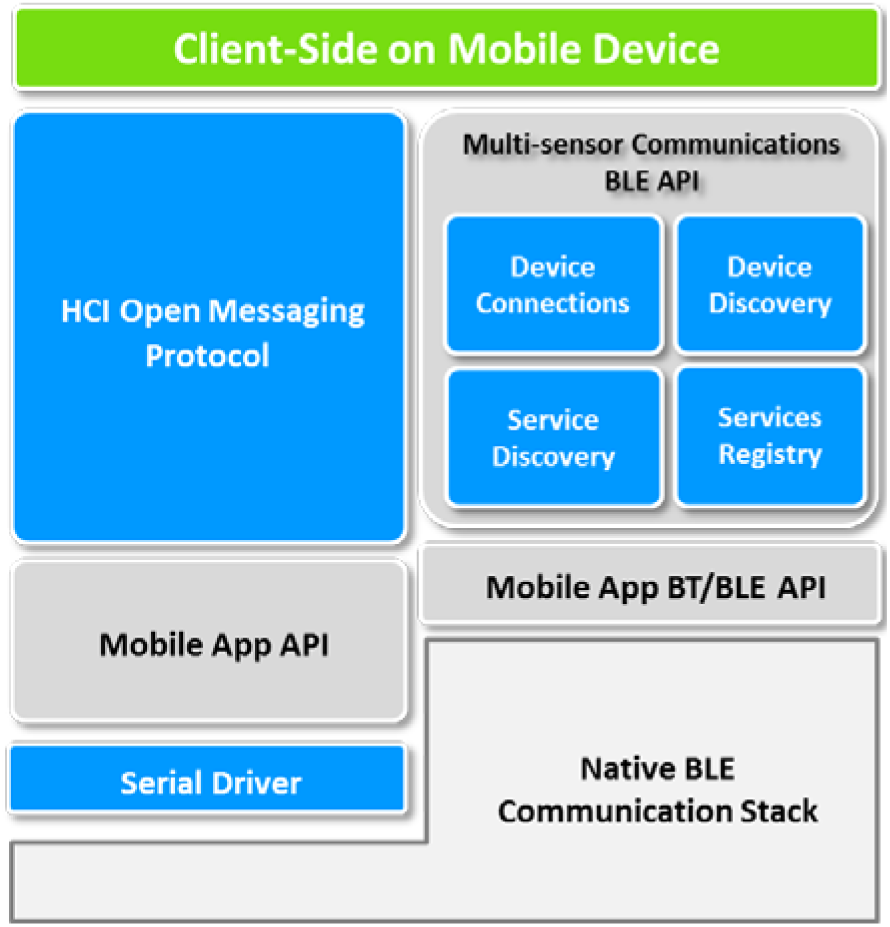
Shows the client-side communication management components (blue) which abstract GATT data exchange and parsing, and abstract the HCI layer into a generic byte level messaging protocol.

**Figure 9.**
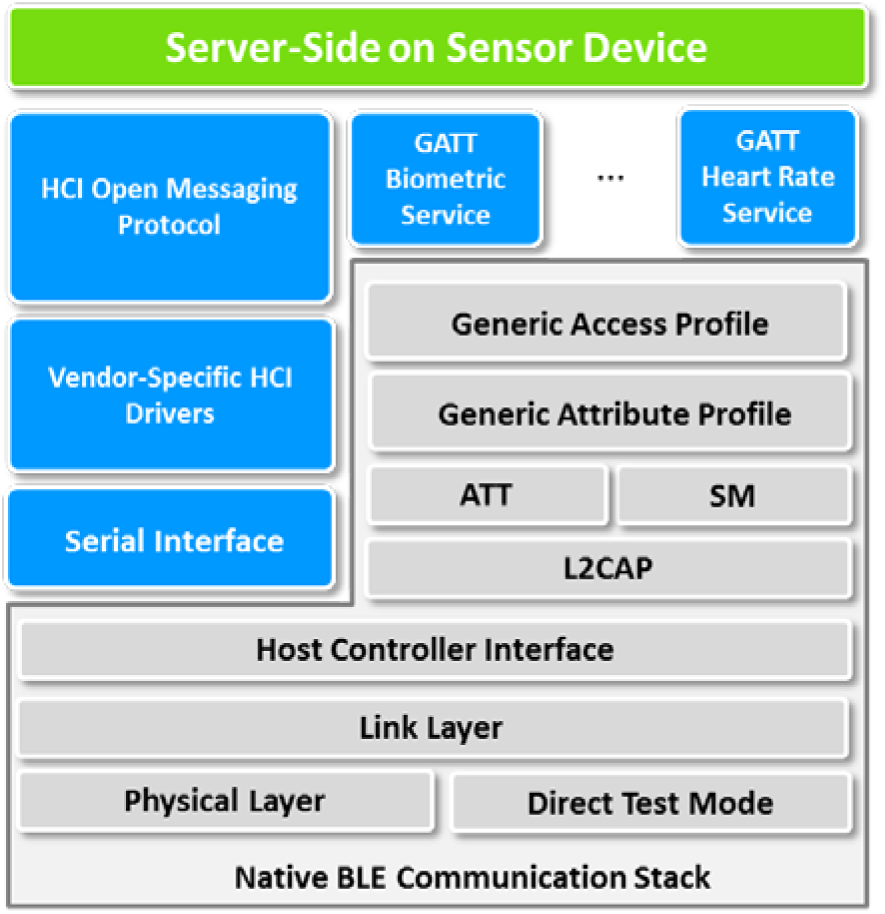
Shows the corresponding sensor-side communication components (blue) for byte level messaging over HCI, or data exchange over ATT/GATT protocols using a new self-describing GATT Biometric service.

On the client-side, in addition to providing components for connections management and device discovery, components for managing and registering GATT services are provided. A mobile application can then either preregister known, certified GATT services which the framework knows how to parse, or by default can extract data contained in a self-describing Generic Biometric Service which is defined in Section II. This allows both commercial sensors using certified Bluetooth SIG services, and prototype sensors using the Generic Biometric Service, to be accessed simultaneously, in a uniform way.

On the server-side, the developer can choose to use either ATT/GATT layers or can use the Host Controller Interface. The framework components at each of these communication levels unify the way data is accessed from a mobile device. As discussed in Section II, the HCI open messaging protocol will provide uniform, communication at byte level and at the GATT layer, a prototype sensor can implement the Generic Biometric Service which allows heterogeneous biodata to be transported and accessed at the mobile device.

## V. FUTURE WORK

A Bluetooth Low Energy based communication framework is proposed with interface layers supporting communications through the ATT and GATT layers or the HCI layer, for capturing data from heterogeneous biosensors, simultaneous and in real-time on a mobile device. These components are intended to be part of a larger open data framework that can provide easy data access, management, and analytics at all layers of the data model discussed in the Introduction. The next steps of this work include a full implementation of the client-server communication components outlined in Section III. Additionally, it may be necessary to support other wireless communication standards. A number of body sensor network frameworks [4, 5, 7] exist which build in support for multiple communication protocols used at the sensor network level. These frameworks will be investigated to see if they can be leveraged or expanded to include the BLE components proposed in this paper.

Communication standards and open sourcing enable the research and commercial communities to make rapid advancements and gain new insights into human health which can aid in providing future knowledge on medical treatments and better health for all. The ultimate goal of this work is to open source the data framework produced for body sensor network application development.

## ACKNOWLEDGMENT

We would like to David Aubin, Erik Metzger for technical feedback and Emily Simons for graphic artwork.

